# Linguistic evolution minimizes the amount of syntactic hierarchy in grammar

**DOI:** 10.1101/2025.07.19.665693

**Authors:** Carlo Y. Meloni, Chundra A. Cathcart, Jessica K. Ivani, Taras Zakharko, Erik J. Ringen, Balthasar Bickel

## Abstract

A core characteristic of language is the capacity for hierarchical syntax, but it remains debated to what extent languages favor hierarchical structure in their grammars, that is, the extent to which they favor hierarchical structures like “When driving, look ahead!” over concatenations like “You drive, you look ahead!”. Through phylogenetic modeling of the grammars of clause combining in three extensive language families (Tupí-Guaraní, Sino-Tibetan, and Indo-European), we find an overall bias towards minimizing hierarchy, with significant cultural variation. Our findings suggest that the computational and representational benefits of hierarchical structure tend not to be universally exploited in clause combinations, likely because they come with additional costs for learning. In consequence, the amount of hierarchy in language might differ from what is found in animal communication in a more graded way than commonly assumed.

---

Non-human primates show remarkable skills in recognizing and producing complex hierarchical structures in experimental tasks [1, 2, 3, 4]. However, such structures seem to be rare in their communication through vocal calls or gestures [5, 6]. This is in stark contrast to the human communication system, language, which is generally thought to exploit hierarchical structure very intensively. But the extent to which this is indeed the case across languages is in fact unknown, and so it remains debated how large and substantial the difference is between animal and human communication, with consequences for our understanding of how language emerged in the hominin lineage.

Languages allow both hierarchical and nonhierarchical structures for expressing what is essentially the same message — in English, for example, we have a choice between (i) hierarchical expressions like “when driving, look ahead”, with an asymmetry between a dependent clause marked by “when” and a main clause that together encode a fixed meaning relation of temporal overlap, and (ii) flat concatenations like “you drive, you look ahead!”, with a symmetry of two main clauses of similar structure that leaves it to context for guessing the intended meaning. Many languages do not leave much choice, however, because their grammars only allow one or the other structure, often depending on the meanings to be expressed [7, 8, 9]. Some languages have been reported to generally disfavor hierarchical structure in their grammars [10, 11, 12, 13], while a phylogenetic study of Indo-European suggests a bias towards hierarchical structures in noun phrases [14]. Given these conflicting results, what, if anything, is the global trend?

One possibility is that languages are shaped by a neuro-cognitive propensity for hierarchical structure in humans, *dendrophilia* [15]. *Dendrophilia is expected to bias languages to maximally exploit the computational power shared with other primates. In computational terms, this power is supra-regular* [1, 3], which removes constraints on branching direction and allows compact representation [16, 15]. For example, in such a system, dependent clauses can be used to add additional information in a sentence (“a big concern [when driving] is micro-sleep”) while keeping the relations local (both “big” and “when driving” are adjacent to “concern” and, at the same time, “concern” remains adjacent to “is” at a higher level in the hierarchy). Compactness and locality are arguably beneficial for (linguistic) thought and creativity [17] as much as for efficient communication [18].

Another possibility is that languages exploit this computational capacity only minimally. Despite the advantages of supra-regular computation, every asymmetry, such as the precise difference between “when driving” and “you drive”, must be learned when languages are transmitted over generations. This favors the simple concatenation of symmetrically structured expressions [19]. Given that utterances are embedded in rich contexts and that “inference is cheap” [20], such concatenations remain fully successful in communication [21, 22]. From this perspective, languages might be expected to minimize the use of hierarchical structures, incorporating more complexity only for special purposes, for example, to distinguish specific meanings [8].

A third possibility is that the evolution of hierarchical structure is essentially random, devoid of direct functional benefits [23]. According to this view, the degree of hierarchy in a grammar would be driven chiefly by idiosyncrasies of language change and perhaps social factors of signaling group differences, schismogenesis [24, 25].

Critical data to test these hypotheses comes from grammars, the structures that need to be computed and learned. We focus specifically on how grammars regulate the combination of clauses. We define hierarchy in clause combination in terms of the presence of structural asymmetries between clauses (“S”), the minimal requirement for a partially ordered set, or, equivalently for current purposes, a directed acyclic graph (*S*_*1*_ →*S*_2_) or a labeled constituent ([*S*_*1*_*S*_1_*S*_2_]) (cf. *Materials and Methods*). We quantify the extent of hierarchy by counting the number of asymmetry features present in a combination of clauses [26, 27]. For example, in the sentence “When driving, look ahead” there are three asymmetry features in terms of the properties by which the first clause differs from the second clause: (1) the presence of “when”, implying the presence of a main clause to which it refers; (2) the absence of a subject expression in the first clause, requiring co-reference with the subject of the second clause; and (3) the absence of finite verb morphology in the first clause, implying inheritance of information from the second clause. By contrast, “You drive, you look ahead” shows none of these asymmetries, hence no reflex of hierarchy in its syntactic structure.

We sampled languages (*N*_lang_ = 59) from three families with highly diverse histories (in terms of literacy, subsistence, environment, and population sizes) and exhaustively coded clause combining constructions for the meaning they express and for whether or not they show an asymmetry across features (*N*_feat_ = 18), e.g. whether or not one of the clauses limits the expression of subject agreement (‘SubjAgrLim’) or its position in linear order (‘OrderLim’) (Figure 1 and *Materials and Methods*, Table 1): Indo-European (IE), Sino-Tibetan (ST), and Tupí-Guaraní (TG).

**Table 1.**
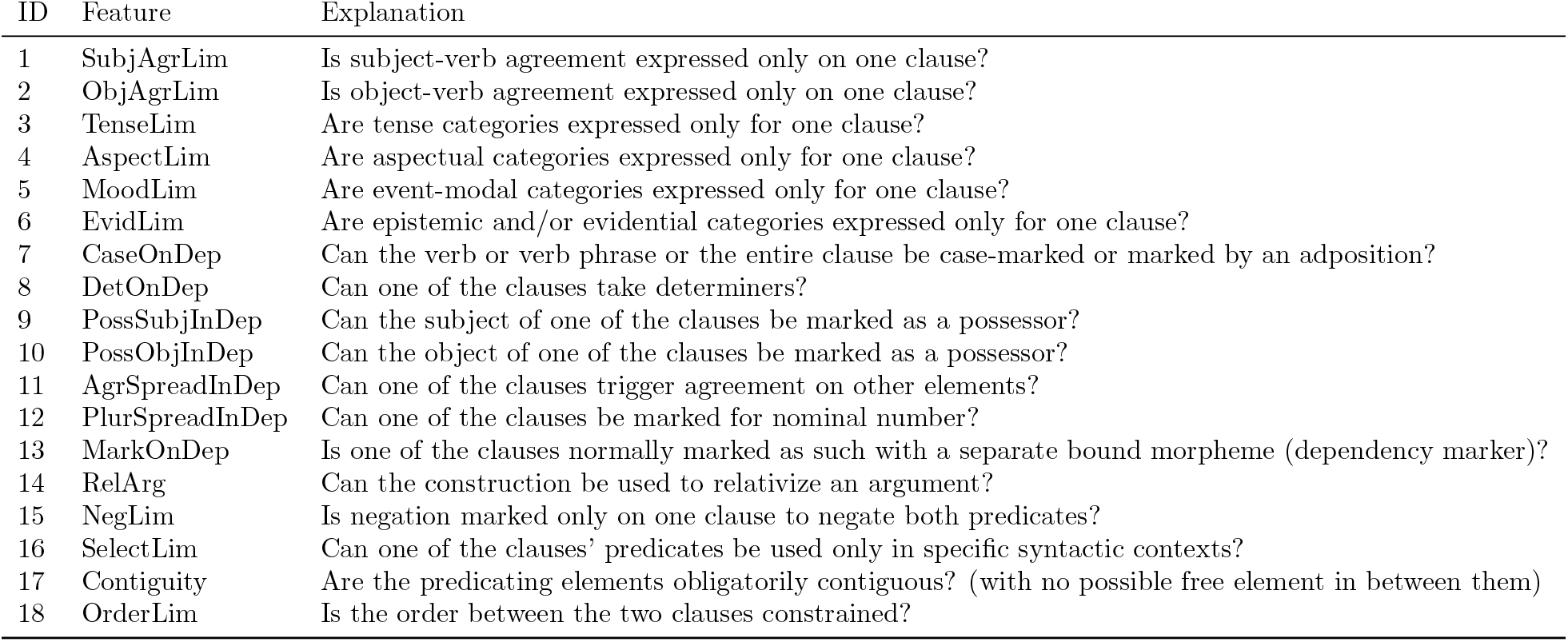
Asymmetry features. See our data file (over the Zenodo repository) for the coding in individual languages.

**Figure 1.**
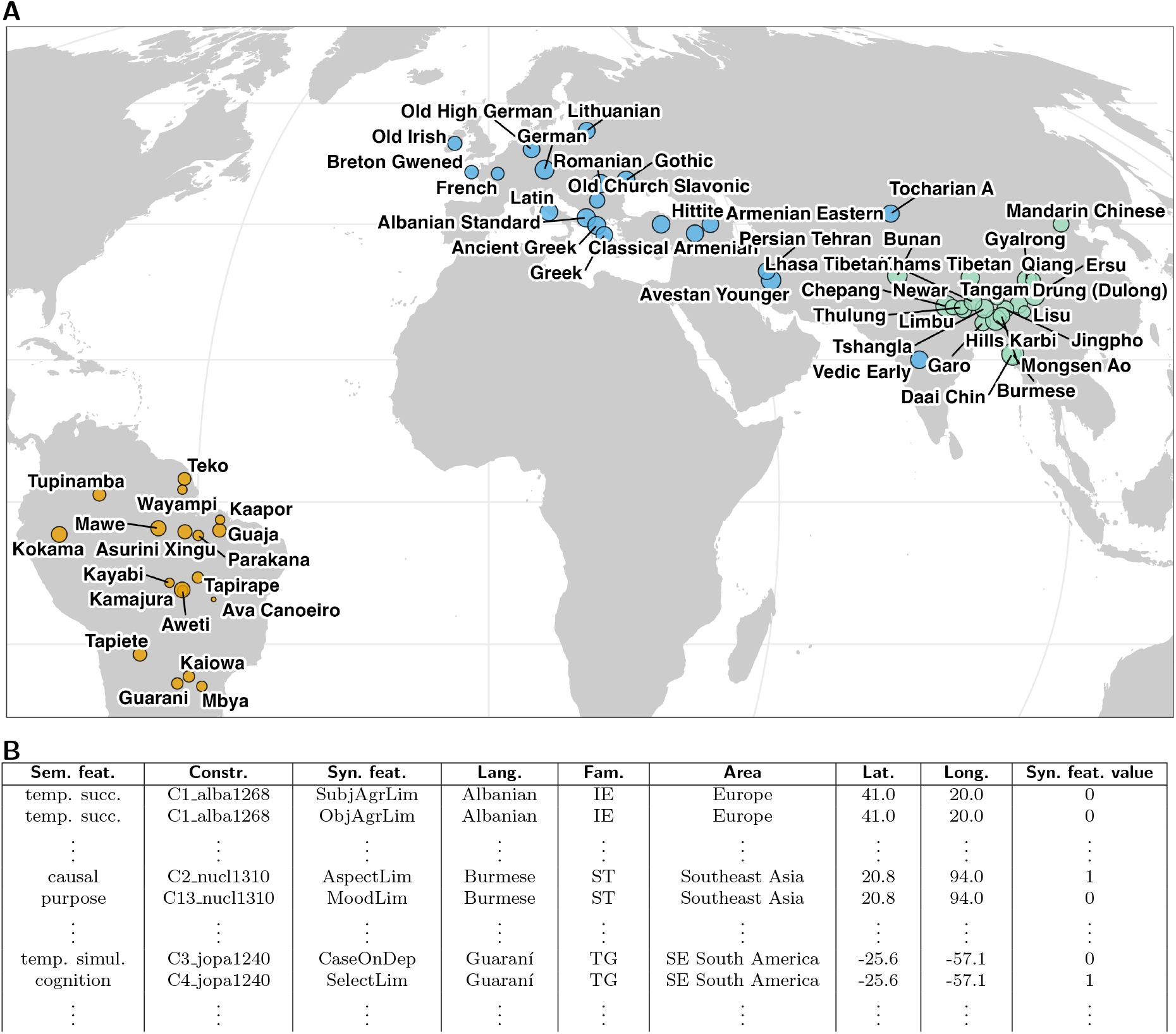
Data on clause combining constructions from 58 languages. **A**: Geographical locations, with dot sizes indicating the number of constructions reported in grammars and colors indicating language families: Indo-European (ie, blue), Sino-Tibetan (st, green), Tupí-Guaraní (tg, yellow). **B**: Constructions were coded for the core meaning relations they cover and whether or not they impose an asymmetry on the clauses with regard to 18 predefined features, exemplified here by limitations on the expression of subject agreement (‘SubjAgrLim’), object agreement (‘ObjAgrLim’), tense (‘TenseLim’), and aspect (‘AspectLim‘) and on the position (‘OrderLim’) of one of the clauses.

## Results

We first selected three exemplars of clause combinations with meaning relations that received scholarship expects to differ in asymmetry [7, 8, 9]: coordination by ‘and’ relations (expected to be least asymmetric), subject relative clauses encoding noun modification (most asymmetric), and constructions expressing ‘because’ relations (varying asymmetry). In each construction, we then modeled the evolutionary dynamics between the presence and absence of each asymmetry feature using separate binary Continuous-Time Markov (CTM) models for each [28, 29].

Figure 2A shows the stationary probability of each asymmetry feature as a measure of long-term trends in evolution (cf. *Materials and Methods*). A meta-analysis of these probabilities reveals an overall trend towards few asymmetries (posterior Pr(Pr(Asymmetry) < .5) > .95 across features; Meta-Analysis I, Supporting Information, Table S1), with only two exceptions where the number of features trending towards more vs. fewer asymmetries are in balance (relative clauses in Sino-Tibetan, median Pr(Asymmetry) = .50, 95%HPDI = [.39, .61], and in Tupí-Guaraní, median Pr(Asymmetry) = .46, 95%HPDI = [.35, .57]). Notably, none of the constructions shows a bias towards asymmetry across all features (Pr(Pr(Asymmetry) > .5) < .5, Supporting Information, Table S1).

**Figure 2.**
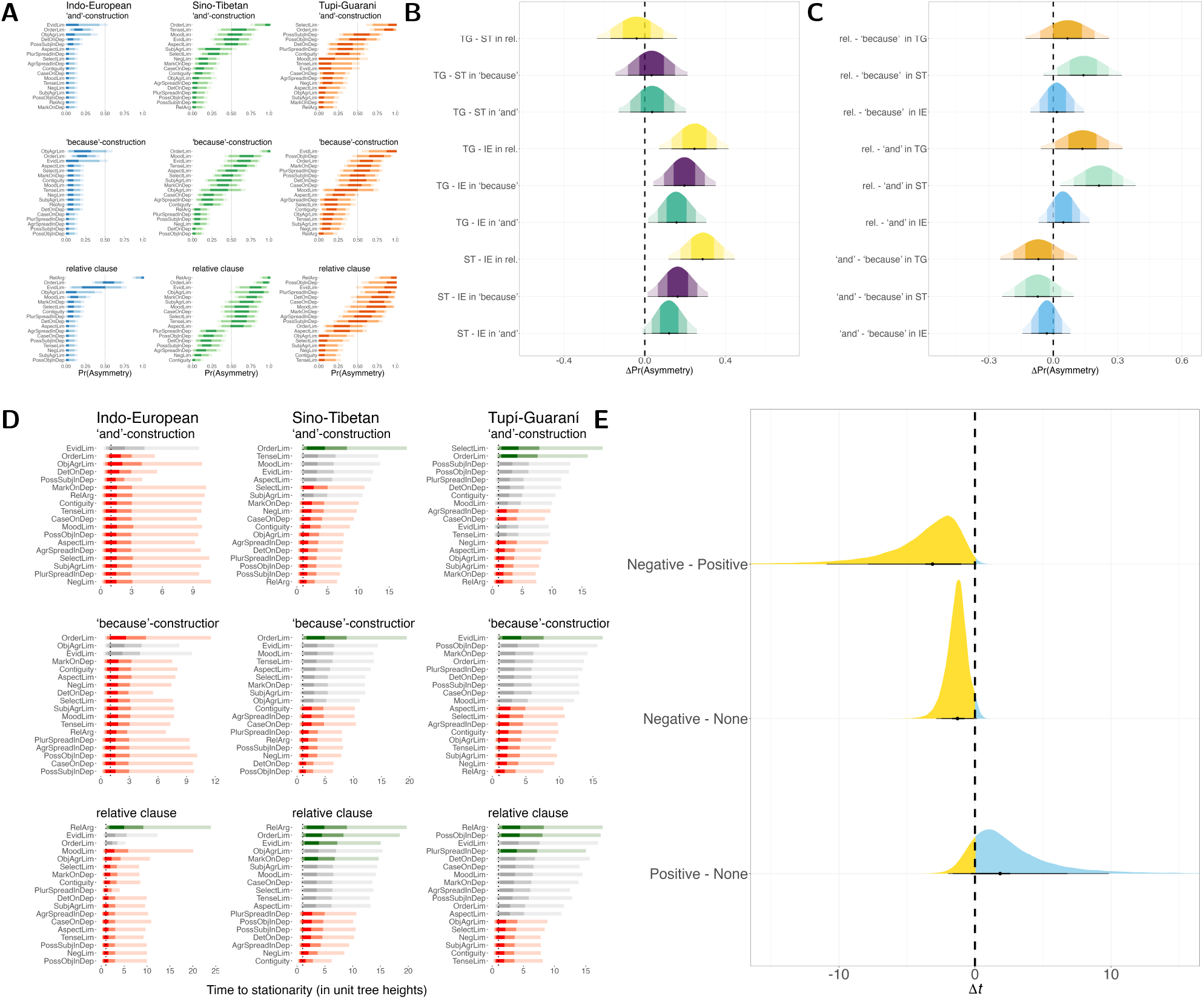
Individual constructions. **A**: Posterior stationary probabilities of the presence of asymmetry features in three selected constructions estimated by Continuous-Time Markov (CTM) models of evolution trend against asymmetry. **B, C**: Average marginal differences in asymmetry probabilities (Δ Pr(Asymmetry)) are stronger between language families (within constructions; **B**) than between constructions (within language families; **C**). **D**: Time required to reach stationarity for each asymmetry feature across families and constructions; the dotted line indicates the time depth of the phylogenies, which we normalized to 1. Red indicates a negative bias, i.e. against asymmetry (Pr(Pr(Asymmetry) < .5) > .95 and green a positive bias, i.e. for asymmetry (Pr(Pr(Asymmetry) > .5) > .95. **E**: Average marginal differences show that features with a negative bias reach stationarity faster (negative Δ*t*) than those with a positive or no bias. Densities in A-D are plotted with 0.5, 0.89, and 0.95 highest posterior density intervals (HPDIs).

At the level of individual features, only few show a bias towards asymmetry (i.e. with Pr(Pr(Asymmetry) >.5) > .95 in Figure 2A). One of them (‘RelArg’) does so trivially because it codes whether or not one of the arguments is relativized, a necessity in relative clause constructions. The others chiefly reflect grammatical constraints on clause order, but this bias is not shared by all constructions and families.

Intriguingly, features with a bias against asymmetry (i.e. with Pr(Pr(Asymmetry) < .5) > .95, colored red in Figure 2D) reach stationarity much faster than features with a bias towards asymmetry (Pr(Δ*t <* 0) = .99) or no bias (Pr(Δ*t <* 0) = .97) (Figure 2E), and do so mostly within the depth of each phylogeny (scaled to unit height in our models; cf. *Materials and Methods*, Meta-Analysis II). By contrast, features with a bias towards asymmetry do not reach stationarity faster than features without a bias (Pr(Δ*t <* 0) = .16). These time estimates suggest that grammars quickly evolve so as to reach stationary probabilities in features that disfavor asymmetry but take much more time (typically beyond the time depth of the phylogenies) when it comes to features that favor asymmetry or show no bias.

Families and constructions vary greatly in the rank order of stationary probabilities per feature (Figure 2A), with no apparent clustering of features per family, construction, or domain of grammar (agreement rules or limitations or tense). Most of this variance stems from a major difference between Indo-European and the other families (Figure 2B, *Materials and Methods*, Meta-Analysis I). Stationary probabilities vary much less between constructions (Figure 2C), with only one difference excluding 0 in its 95% HPDI (the difference between ‘and’-constructions and subject relative clauses in Sino-Tibetan). This suggests that asymmetries depend less on the relations they encode (here, ‘and’ vs. ‘because’ vs. relative clauses) than is traditionally assumed.

Given that asymmetry features do not differ much across constructions and meanings, we next tested the global trend towards low asymmetry in a multi-level phylogenetic model that aggregates the information from all constructions and meanings (i.e., not only the selected three in the CTM model). To this end, we fitted a model that assumes the logit-transformed presence of asymmetry to evolve as a continuous trait and that accommodates extra variation between features, languages, constructions, and encoded meaning relations (cf. *Materials and Methods*).

Leave-one-out cross-validation suggests that the asymmetries evolve according to an Ornstein-Uhlenbeck process [30, 31, 32] (IE: difference in expected log-pointwise predictive densities Δ_ELPD_ = 35.4, SE = 7.4, stacking weight *w* = 100%; ST: Δ_ELPD_ = 59.1, SE = 10.0, *w* = 100%; TG: Δ_ELPD_ = 61.5, SE = 6.1, *w* =100%). The Ornstein-Uhlenbeck process differs from a Brownian motion process by adding to basic stochastic fluctuation *η* a stationary value *θ* (here, the logit-transformed binomial counts of asymmetries) to which evolution is attracted with selection strength *α*. We report selection strength in terms of what is known as phylogenetic half-lives 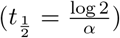 [33]. This is the expected time it takes for the attraction to *θ* to outweigh the influence from the ancestral state [34, 33]. (To further help interpretation of how the three parameters shape evolution, Supporting Information Figure S2 shows simulations of the evolutionary dynamic arising from the model estimates).

Figure 3 shows that the stationary probabilities of asymmetries are strikingly low (IE: posterior median logit^™1^*θ* = .2, 95%HPDI =[.09,.35], Pr(logit^™1^*θ <* .5) = .99; ST: logit^™1^*θ* = .27, 95%HPDI = [.08, .52],Pr(logit^™1^*θ <* .5) = .95; TG: logit^™1^*θ* = .31, 95%HPDI = [.15, .5], Pr(logit^™1^*θ <* .5) = .96), consistent with the trend emerging from the CTM models of individual constructions. Asymmetry features vary around the overall trend to different extents in each family (Figure 3B), again similar to the CTM model results.

**Figure 3.**
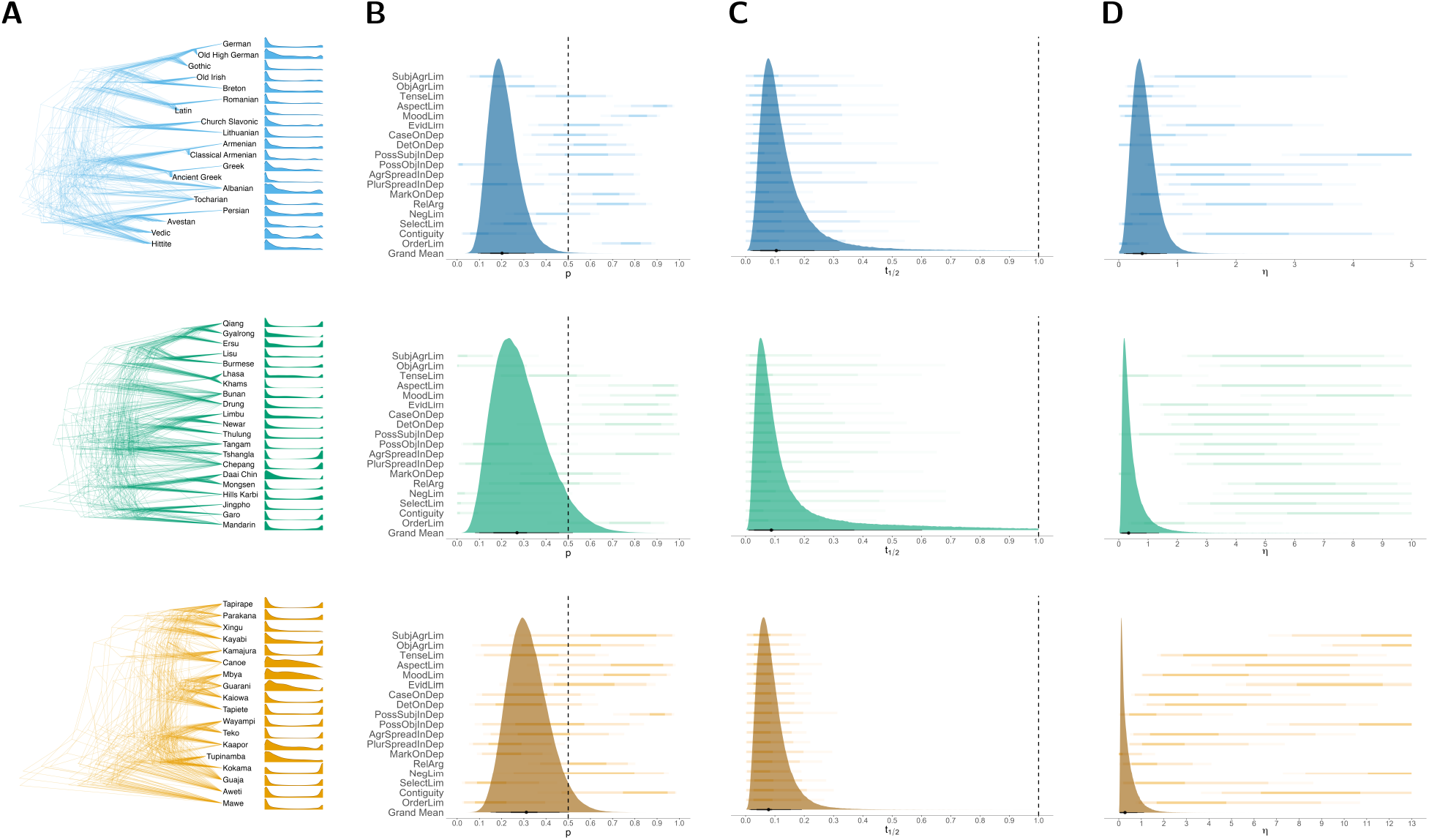
Evolutionary dynamics across constructions according to a multilevel phylogenetic model in which the logit-transformed presence of asymmetry features evolves according to an Ornstein-Uhlenbeck process. **A**. Posterior predicted probabilities of asymmetries per language mapped to posterior draws from phylogenies. **B**: Posterior stationary probabilities per asymmetry features (ordered as in Table 1) and grand mean across features (logit^™1^*θ*). **C**: Posterior estimates of the phylogenetic half-lives 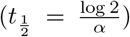 as an indicator of selection strength. (All times are at unit scale, with 1 representing the time depth of each sampled phylogeny.) **D**: Posterior estimates of the stochastic fluctuation over time (*η*).

Selection is strong in all families (Figure 3C) since the 95% credible interval of the phylogenetic halflives excludes the time depth of each language tree (scaled to unit in the models; IE: posterior median 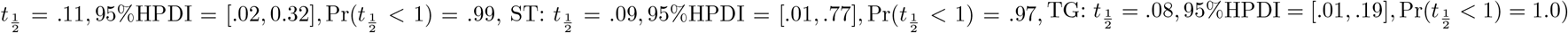. Unlike stationary probabilities, features do not vary much in half-lives, but families differ in their posterior spread, chiefly as a result of sample size variation.Despite this, the half-life estimates consistently show that enough time has elapsed for evolution to have converged on the stationary optimum of *θ* and thereby on the expected long-term probabilities of asymmetry across features. This is again consistent with the trend observed for individual constructions, where features with a bias against asymmetry mostly reached stationarity within the time depth of each family, much faster than features with a bias towards asymmetry or no bias (Figure 2D).

Stochastic variation (*η*) is also roughly similar across families (Figure 3D), despite notable differences in the posterior spreads (ST: posterior median *η* = .36, 95%HPDI = [.05, 1.37]) than the other two families (IE: *η* = .39, 95%HPDI = [.09, .82], TG: *η* = .28, 95%HPDI = [.03, 1.1]). Like in the case of stationary probabilities, stochastic variation affects individual features in different ways across families.

## Discussion

Our results speak against a global bias for hierarchical structure in clause combinations. Evolutionary models of individual features per construction and of all features aggregated across constructions suggest an overall bias against asymmetries and therefore against hierarchical structure. Across models, our results furthermore suggest strong evolutionary pressure to return to this bias, so that languages evolve relatively fast into a stationary optimum with low asymmetries, mostly within the time depth of reconstructed language families. Together, our results suggest that the evolution of hierarchy in clause combining is strongly shaped by constraints on learning, favoring simpler structures despite the potential benefits offered by hierarchical structure.

At the same time, we note considerable variation in individual features across language families, suggesting a fair amount of cultural variation. This finding is consistent with the notion that a key property of the human language faculty is a strong drive towards diversity in response to social grouping [25]. The variation seems largely independent of meaning and construction types, consistent with earlier findings that challenged the notion of universally recurrent patterns in clause combinations [26].

Clearly, these propositions need further testing. One limitation of the present study is that we only examined grammar data. It might be that hierarchical structures are specifically exploited in usage, although most reports from spontaneous spoken data point to the other direction [35, 36] and we found no difference in a large-scale study of 59 languages [37]. A second limitation is that we only modeled data from clause combining. It is unclear whether our findings generalize to other aspects of language. Noun phrase structures favor some amount of hierarchy in the evolution of Indo-European [14], but the evidence for this is limited to one language family, and it rests on the sheer presence of hierarchy, not quantifying its extent in terms of asymmetries that need to be learned.

In conclusion, our findings invite a re-appreciation of the contrast between animal and human communication in terms of hierarchical structure. The extent to which the shared capacity for such structure is exploited in human communication might have been overestimated. Recent findings from animal behavior suggest that, conversely, the extent to which nonhuman primates can exploit hierarchical structure in call combinations might have been under-estimated [38, 39, 40, 41].

## Materials and Methods

### Data and coding

We sampled a total number of 696 constructions (Indo-European: 285; Sino-Tibetan: 300; Tupí-Guaraní: 111), defined by unique combinations of asymmetry features and meaning relations following an established schema [27]. All the data is accessible in the [Zenodo repository](https://doi.org/10.5281/zenodo.16105001).

Our operationalization of hierarchy is based on the concept that syntactic structure corresponds to partially ordered sets with a non-associative operator for the formation of the sets. This results in trees (in the sense of directed acyclic graphs or labeled bracket structures) that are sensitive to their generative processes, capturing the notions of compositionality and structure-dependence in linguistics [42]. Importantly, the evidence for dependency arcs or labeled bracketing comes from specific syntactic and semantic asymmetry features imposed by the grammar of each language (Table 1). We quantify hierarchy by the number of such features in individual constructions and across all constructions in a language. This approach is independent of how hierarchical structure is formally expressed in linguistic theories [43], whether through dependency graphs, nested sets, or constituent structures. Theories differ in how linear order is treated; here, we take constraints on linear order as distinct evidence for asymmetry and therefore hierarchical structure, differentiating the two clauses in a combination into one that must occur first and one that must occur second.

For quantifying hierarchy in individual constructions, we selected exemplars expressing ‘and’-relations, ‘because’relations and noun modification relations relativized on subjects (‘the computer that crashed’). Whenever multiple constructions express the same relation, we choose the one reported as most frequently used and/or most semantically general (i.e., the one with fewer constraints on its usage). If the usage of a construction was not documented in the grammar of a language, or if the constructions did not differ in the relations they express, but showed differences in their asymmetry features (e.g., one construction had a value of 1 while the other had 0), we modeled the relevant language with an uncertain tip state in the relevant feature (Pr(Asymmetry) = .5).

### Continuous Time Markov Model

We used the Bayesian framework of `Stan` [44] to estimate the gain (*α*) and loss rate (*β*) of each asymmetry feature, with the likelihood function computed through the application of the pruning algorithm [45]. We assigned non-informative Gamma(1, 1) priors to both rate parameters. The model was executed on a subset of 100 trees derived from phylogenies of each language family [46, 47, 48], each scaled to unit height. We computed the stationary probabilities for the presence of an asymmetry feature as 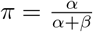.

We performed two meta-analyses on the CTM estimates. In Meta-Analysis I we regressed stationary probability estimates on families and constructions, assuming a Beta likelihood and the flat priors given by default in the `brms`[50] interface to `Stan`. In Meta-Analysis II, we first estimated the time it takes for each CTM to reach stationarity by minimizing the Kullback–Leibler divergence between the time-evolved distribution *π*_*t*_ and the true stationary distribution at infinity 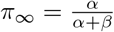 (*Supporting Information*, subsection *Meta-analysis II*). We then regressed these time estimates on whether a feature shows a bias against or towards asymmetry, i.e., Pr(Asymmetry) < .5) > .95 vs. Pr(Asymmetry) > .5) > .95, or no bias, allowing for variation by family and construction and assuming a log-Normal likelihood. To help with convergence, we set a *N* (0, 5) prior on the bias predictor and an Exp(1) prior on the varying intercepts and slopes. In both metaanalyses, we report the average marginal effects across predictors and groups using the `marginaleffects` package [51]. See *Supporting Information*, section *Models* for further specifications.

### Continuous-trait model

We implemented the Ornstein-Uhlenbeck (OU) and Brownian motion (BM) models as an intercept-only phylogenetic regression model allowing for group variation (‘random effects’). The outcome variable is modeled using a Bernoulli likelihood with a logit (log-odds) link:

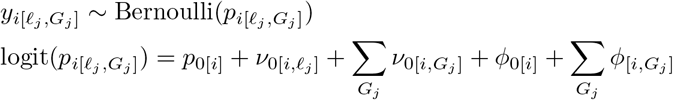

Here, each observation’s log-odds is determined by a global intercept *p*_0[*i*]_, language-level deviations 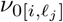, and group-level deviations across asymmetry features, constructions, and semantic relations (three types of *G*_*j*_), each modeled as normally distributed random effects, with Σ representing the variance-covariance matrix of the relevant grouping.

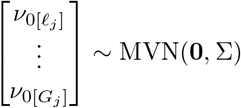

Phylogenetic effects are introduced through Cholesky-decomposed OU covariance matrices, affecting both the global intercept and group-level components:

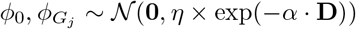

where **D** is the pairwise phylogenetic distance matrix between languages; *η* controls the stationary variance, and *α* determines the strength of attraction under the OU process. These parameters vary by group (features and semantic relations), allowing the model to capture structured correlations induced by evolutionary history. Under the OU process as a stochastic differential equation, 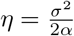.

We placed weakly regularizing priors (*N* (0, 1)) on all model parameters. The model was applied to a sample of 50 trees drawn from the same phylogenies used in the CTM model, each scaled to unit height. Full model details are provided in the *Supplementary Information*, subsection *OU and BM Models*.

For model comparison, we approximated the out-of-sample predictive fit with Pareto-smoothed importance sampling from the posterior [52]. We report model differences in terms of expected log-pointwise predictive densities (ELPD) and the weights allocated to each model when they are stacked to jointly predict out-of-sample data points in a leave-one-out setting [53].

## Acknowledgments

We are grateful to Mathias Jenny (Sino-Tibetan), Rik van Gijn (Tupí-Guaraní), and Paul Widmer (Indo-European) for their help with data collection and to Lothar Sebastian Krapp for advice on the mathematical definitions of hierarchy.

## Contributions

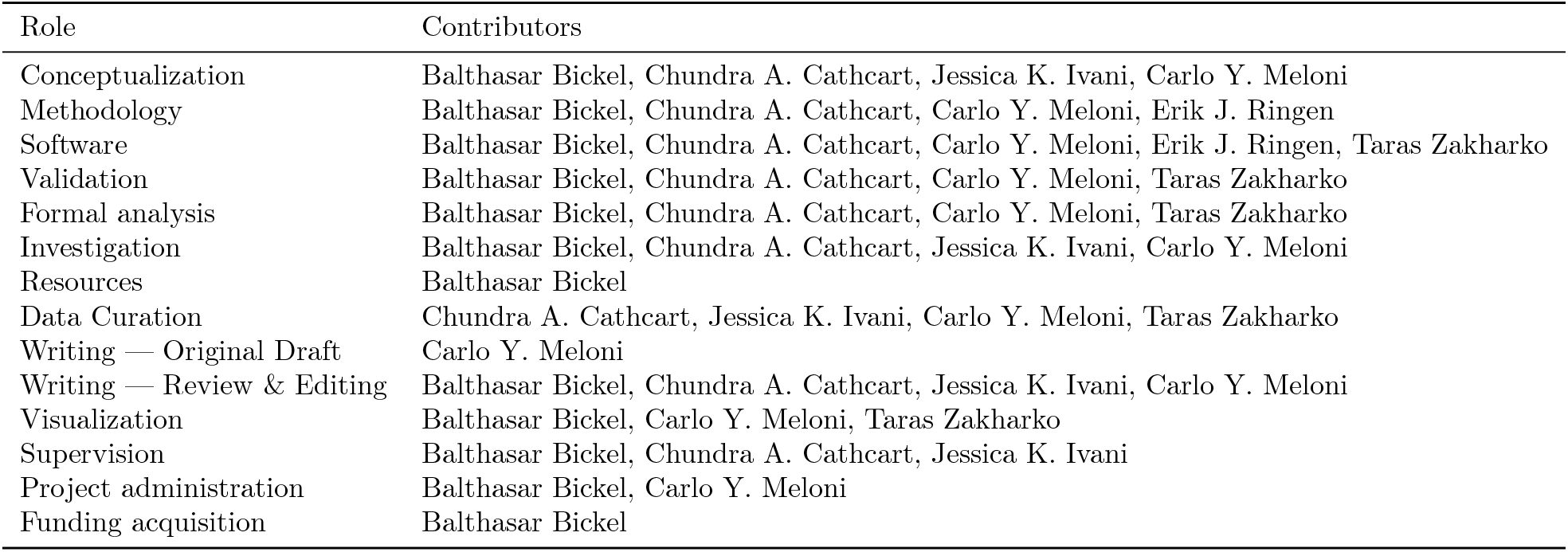

## Supplementary Information

### Data

#### Construction sample

A total of 59 languages are included in our sample, representing three families: Indo-European (IE), Sino-Tibetan (ST), and Tupí-Guaraní (TG).

The total number of constructions sampled was 696, distributed as follows: 285 in IE, 300 in ST, and 111 in TG. The average number of constructions per language was 14.2 for IE, 14.3 ST, and 6.17 for TG. The standard deviation of constructions per language was 3.49 for IE, 4.81 for ST, and 3.28 for TG. Figure S1 illustrates the distribution of constructions across the languages.

A syntactic structure was defined as a separate construction if it differed from a similar syntactic construction in at least one type of asymmetry (see next subsection).

#### Asymmetry types

The sampled constructions were analyzed by decomposing them into an array of 18 distinct asymmetry features, each capturing different aspects of the syntactic relationship between the clauses that compose the constructions. They are based on the framework provided by [27] and are structured as follows:

- IDs 1-6 pertain to the agreement and tense-aspect-mood (TAM) properties of the dependent event-denoting unit (EDU).
- IDs 7-12 address the degree of nominalization within the construction.
- ID 13 concerns the nature of syndesis between the construction clauses.
- ID 14 specifically deals with relativization.
- IDs 15-18 encompass general properties of clause independence, such as negation scope and clause order.

In addition to the asymmetry features listed above, 20 semantic features capture the meanings that can be encoded by a clause combining construction and are used as random effects in our model.

### Models

#### CTM model

For the first analysis, we implement a Continuous Time Markov model (ctm) for reconstructing the gain and loss rate (*α* and *β* respectively) of each of the 18 asymmetry features for three types of constructions (‘and’, ‘because’ and relative clause constructions), along a phylogenetic tree. The likelihood is computed using Felsenstein’s pruning algorithm [54].

For a given branch length *z*, the transient probability matrix *P* (*z*) for transitioning between states is given by:

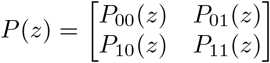

where:

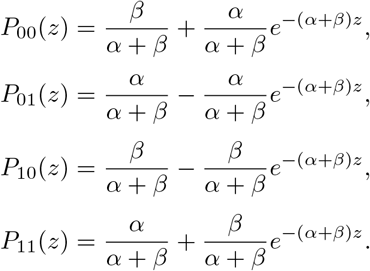

The likelihood calculation via Felsenstein’s pruning algorithm can be formulated as follows: Let *λ*_*n,d*_ represent the log-likelihood of state *d* at node *n*. The likelihood propagation follows:

1. For each tip *t*, initialize:

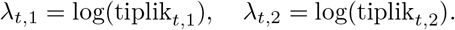
2. For each internal node *n*:

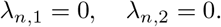
3. For each branch *b* (from child *c* to parent *p*):

- Compute the transition probability matrix *P* (*z*_*b*_).
- Update the log-likelihood at the parent node:

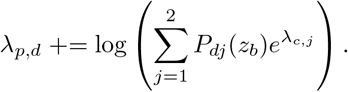

4. Compute the total likelihood at the root node (***π*** represents the prior state distribution at the root of the tree, which we set uniformly to *pi*_1_ = *pi*_2_ = .5):

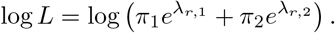

The transition rates *α* and *β* are assumed to follow gamma priors:

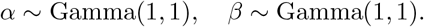

The posterior distribution of *α* and *β* is obtained by maximizing the log posterior:

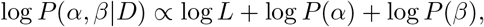

where *L* is the likelihood computed using Felsenstein’s pruning algorithm.

The model was fitted on four parallel chains, each comprising 5000 iterations, with the initial 2500 iterations considered burn-in, and was executed on a subset of 100 trees derived from phylogenies of each language family.

#### Meta-analysis I

We conducted a Bayesian meta-analysis of the stationary probabilities 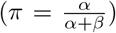 inferred from the CTM posterior distributions using the `brms` package [50]. The probabilities represent the stationary gain of a change process and are bounded between 0 and 1, making the Beta distribution a natural choice.

Let *p*_*ijk*_ denote the stationary probability for the *i*-th sample (*i* = 1, …, 1000), construction type *j*(*j* = 1, …, 3), and family *k* (*k* = 1, …, 3). We model:

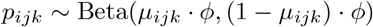

where:

- *µ*_*ijk*_ is the mean of the Beta distribution and is modeled as a function of the predictors.
- *ϕ* is a precision parameter controlling variance.

The mean *µ*_*ijk*_ is linked to the predictors via a logit function:

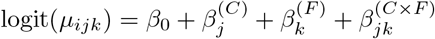

with:

- *β*_0_ the intercept,
- 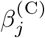 the effect of the *j*-th construction type,
- 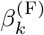 the effect of the *k*-th family,
- 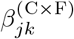 the interaction term between construction and family.

All predictors are categorical, with three levels each (i.e., *N*_Construction_ = 3, *N*_Family_ = 3). The model estimates the parameters using weakly informative priors set by default in`brms`.

#### Meta-analysis II

We modeled the time required for each syntactic feature to reach its stationary distribution using a Bayesian hierarchical regression model implemented in the `brms` package. The outcome variable is the estimated time to stationarity *T*_*ijkl*_ (in units of tree height) for the *i*-th posterior sample of a specific asymmetry feature *l* in a given construction *j* and language family *k*, conditioned on the inferred direction of bias (Bias_*ijkl*_ ∈ positive, none, negative).

#### Computation of Time to Stationarity

The time to stationarity *T* is computed numerically for each posterior draw of the CTM parameters (*α, β*) using a binary search on the Kullback–Leibler (KL) divergence between the time-evolved distribution *π*_*t*_ and the stationary distribution *π*_*∞*_:

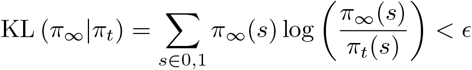

The threshold ϵ is set to 0.001. The process continues until the KL divergence between the evolved and stationary distribution drops below this value, yielding the estimated *T* for that draw.

#### Statistical model

Let *T*_*ijkl*_ be the time to stationarity for the *i*-th sample (draw) of feature *l*, construction *j*, and family *k*. The model is specified as:

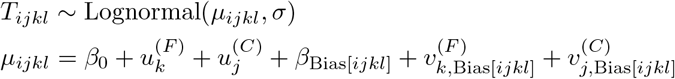

where:

- *β*_0_ is the intercept.
- 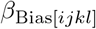 is the fixed effect of **Bias** towards asymmetry (positive, none, negative).
- 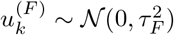 is the random intercept for **Family** *k*.
- 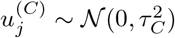 is the random intercept for **Construction** *j*.
- 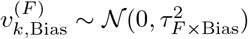 is the random slope of **Bias** within **Family**.
- 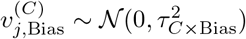 is the random slope of **Bias** within **Construction**.

The model allows for varying intercepts and slopes across both family and construction groupings, capturing heterogeneity in how bias affects the time to stationarity.

The priors used in the model are weakly informative:

- *β* ∼ *N* (0, 5) (fixed effects)
- *σ* ∼ Exponential(1) (residual standard deviation of the log-normal distribution)
- *τ* ∼ Exponential(1) (standard deviations of group-level effects)

Posterior sampling was performed using a single chain of 5000 iterations with `adapt_delta` = 0.999 and `max treedepth` = 15, based on 1000 posterior samples of the CTM parameters (randomly selected from the full posterior sample space).

#### OU and BM Models

The model describes the occurrence of asymmetry features across languages. We model the binary outcome *y*_*i*_ ∈ {0, 1} representing the presence of an asymmetry feature for each observation *i* in our dataset.

Let *y*_*i*_ ∼Bernoulli(*p*_*i*_), where the success probability *p*_*i*_ is defined through a logit transformation of multiple additive components:

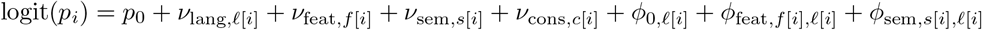

where:

- *p*_0_: global intercept on the logit scale.
- *ν*_lang,*ℓ*_ ∼ *N* (0, *σ*_lang_): language-level random intercepts.
- *ν*_feat,*f*_ ∼ *N* (0, *σ*_feat_): feature-level random intercepts.
- *ν*_sem,*s*_ ∼ *N* (0, *σ*_sem_): semantic-group-level random intercepts.
- *ν*_cons,*c*_ ∼ *N* (0, *σ*_cons_): construction-level random intercepts.

All random effects are parameterized as standard normal samples scaled by a standard deviation parameter, enhancing computational efficiency in `Stan`, while being mathematically equivalent to *N* (0, *σ*).

The model accounts for three phylogenetic components based on the patristic distance matrix **D** between languages, where GP is a Gaussian process (explained below):

- Global phylogenetic effect *ϕ*_0_ ∼ GP_OU_(*η*_0_, *α*_0_).
- Feature-specific phylogenetic effect *ϕ*_feat,*f*_ ∼ GP_OU_(*η*_*f*_, *α*_*f*_).
- Semantic-specific phylogenetic effect *ϕ*_sem,*s*_ ∼ GP_OU_(*η*_*s*_, *α*_*s*_).

The phylogenetic components are modeled using a stationary Ornstein-Uhlenbeck (OU) model, defined by the stochastic differential equation (SDE):

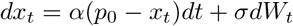

Here, *α* controls the strength of mean reversion, *σ* is the scale of the Wiener process *W*_*t*_. When stationary, the OU process can be represented as a Gaussian Process (GP) covariance function. Given the patristic distance matrix **D**, the phylogenetic covariance *K*_OU_ between two languages *j* and *j*^*′*^ is:

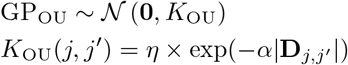

Where *η* represents the total phylogenetic variance, defined as 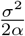, and is analogous to *σ* in the SDE formulation of the OU process [55]. The optimum *p*_0_ is external to the covariance function, consistent with the assumption that the GP has a stationary mean of 0. Scaling **D** by its maximum value facilitates setting priors.

As *α*→0, the OU model approximates a pure-drift Brownian motion (BM) process with stationary increments:

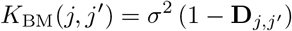

The OU parameters vary hierarchically:

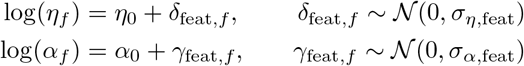

with analogous equations for the semantic groups.

All non-centered parameters are sampled from standard normal distributions and scaled:

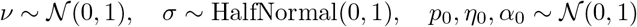

For each observation *i*, we model *y*_*i*_ ∼ Bernoulli(*p*_*i*_) and simulate posterior predictive draws 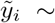 Bernoulli(*p*_*i*_), with associated log-likelihood 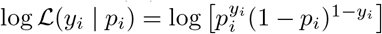

The model was fitted on four parallel chains, each comprising 5000 iterations, with the initial 2500 iterations considered burn-in, and was executed on a subset of 50 trees derived from phylogenies of each language family.

---

**Figure S1.**
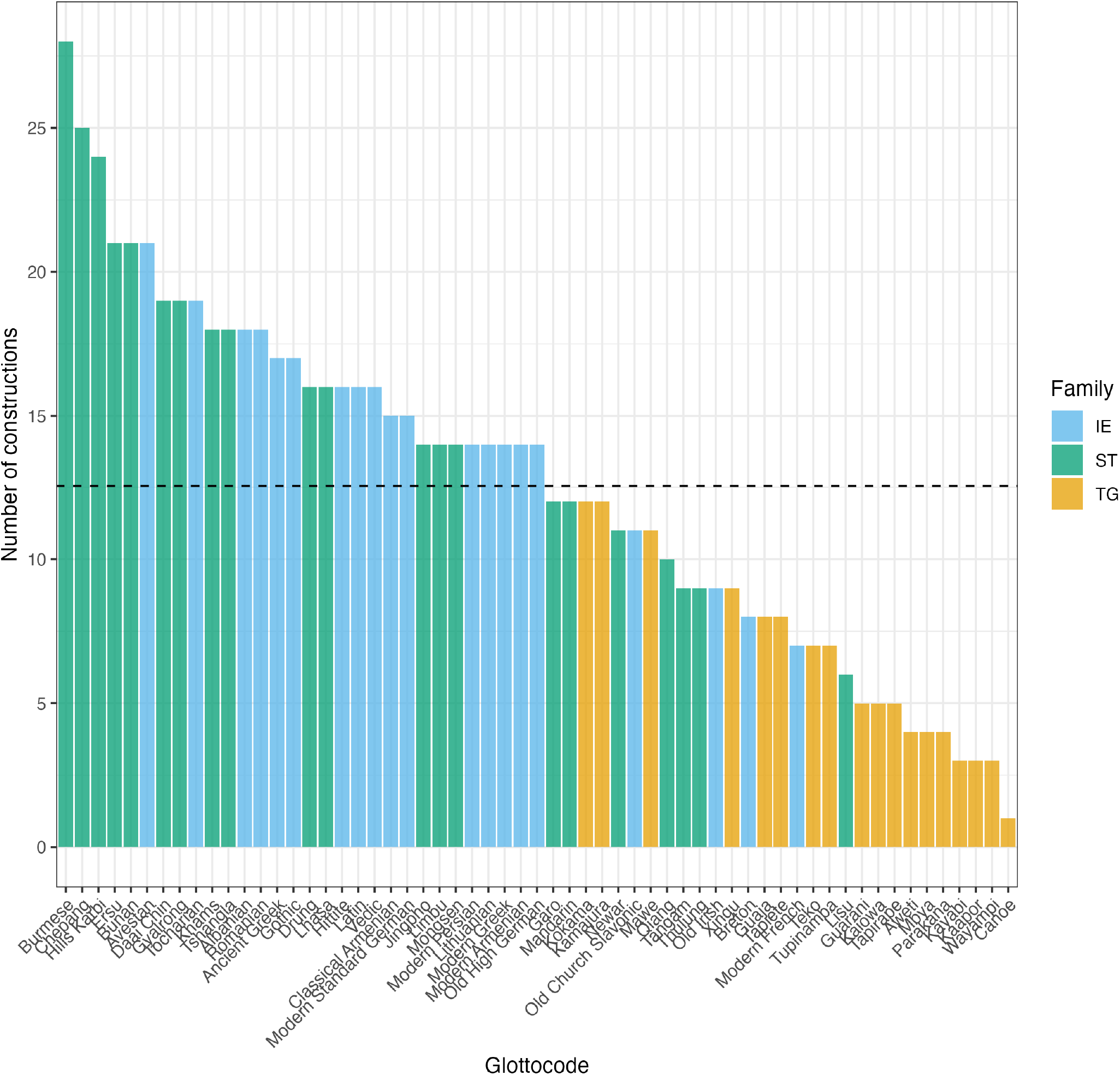
Number of constructions per language. The horizontal dashed line represents the mean construction number.

**Figure S2.**
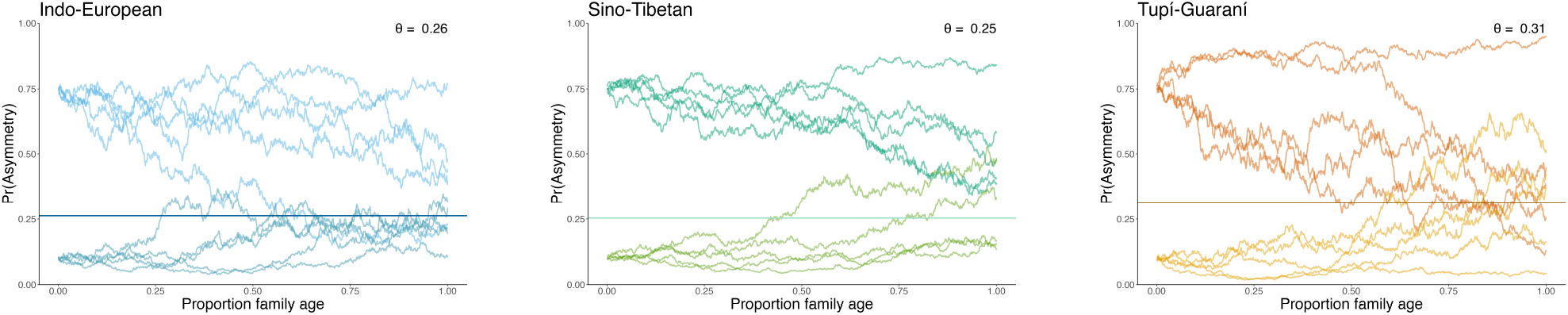
Simulations of the Ornstein-Uhlenbeck process employed for modeling asymmetric syntactic construction change dynamics. The ou process is characterized by mean reversion, implying that over time, the process converges back to the mean or optimal (*θ*) value, although temporary deviations are possible. Our findings suggest a propensity against asymmetric values, with an estimated *θ* ranging between 0.25 and 0.31, contingent upon the language family. The figure illustrates that the initial state (i.e., the initial ratio of asymmetric values) does not exert an impact on the long-term process. Each line in the plots represents a simulated lineage from a language family.

**Figure S3.**
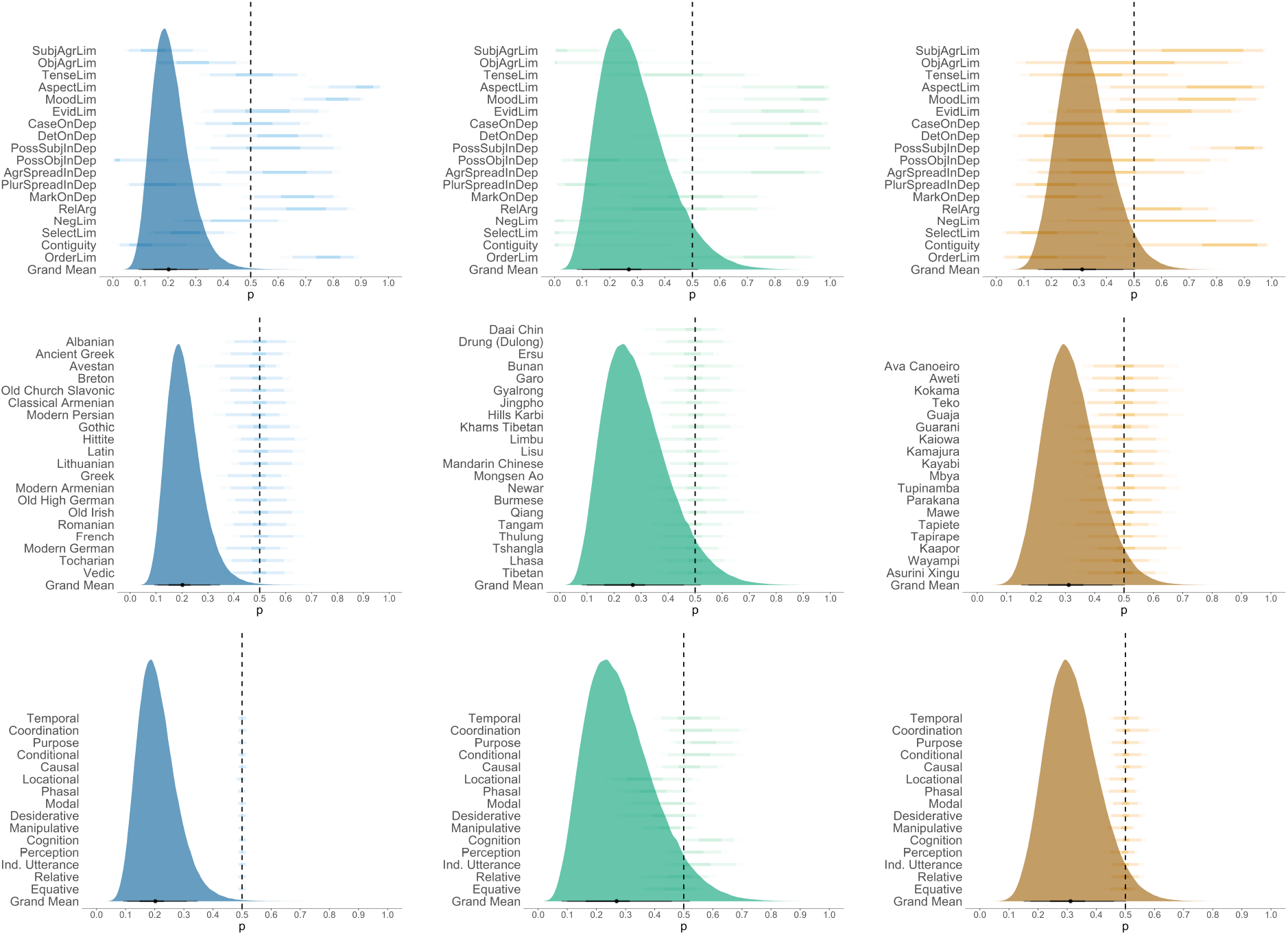
Posterior stationary probabilities (logit^™1^*θ*) for different random effects and the grand mean across them. The columns represent different language families: Indo-European, Sino-Tibetan, and Tupí-Guaraní. The rows correspond to distinct random effects: (i) asymmetry features, (ii) language-specific variation, and semantic features.

**Figure S4.**
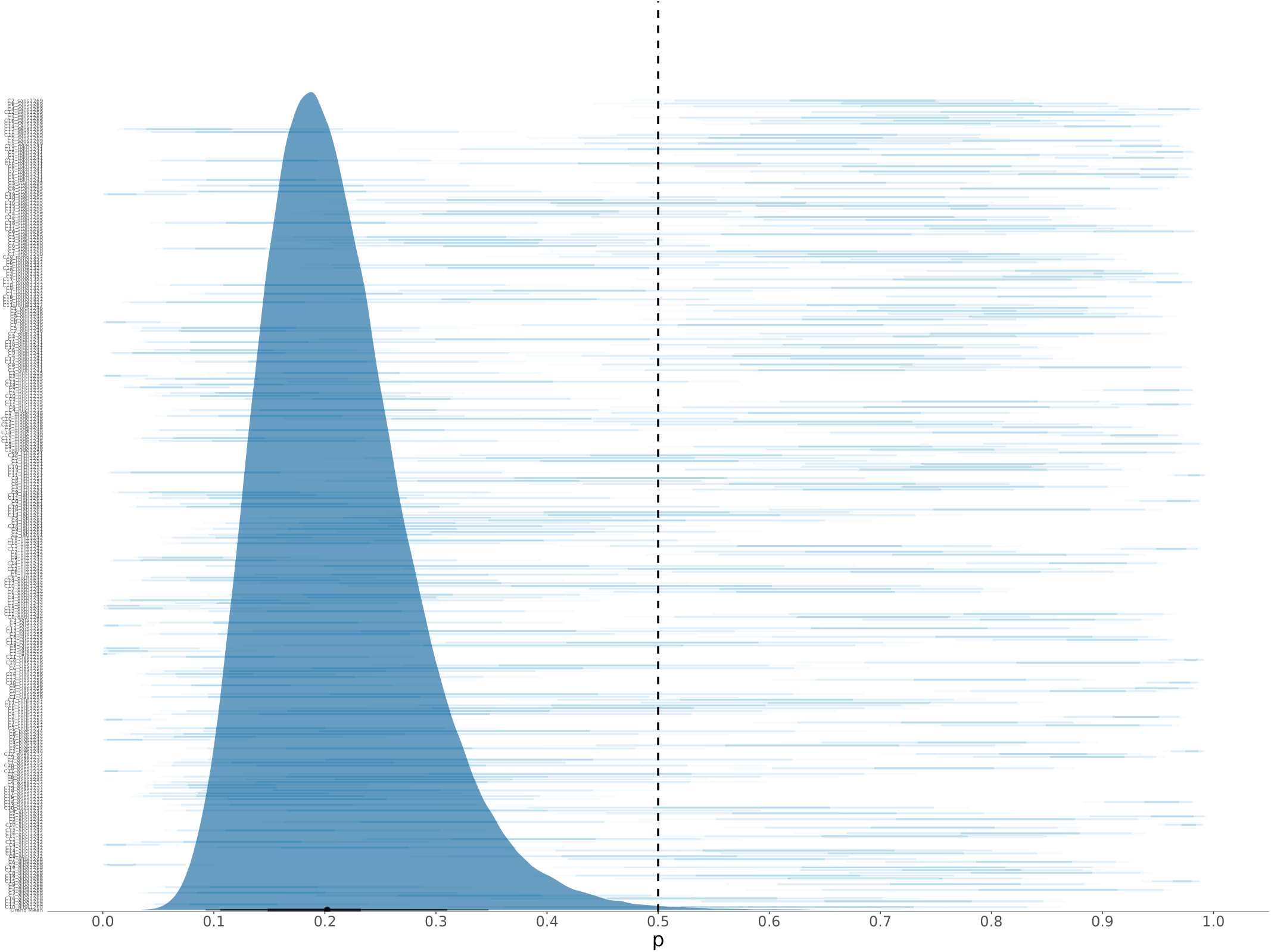
Posterior stationary probabilities (logit^™1^*θ*) for specific syntactic constructions random effects, and the grand mean across them (IE).

**Figure S5.**
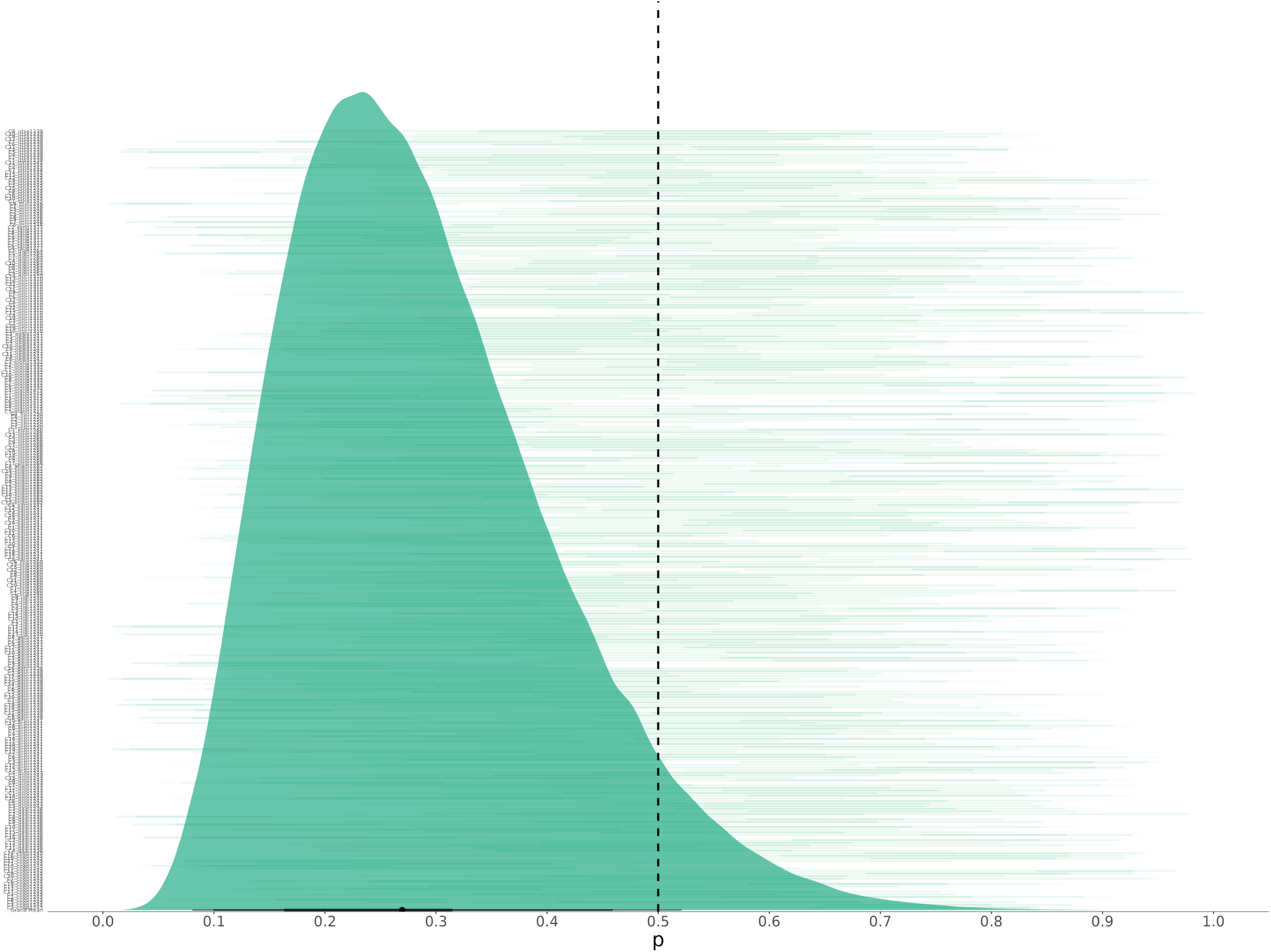
Posterior stationary probabilities (logit^™1^*θ*) for specific syntactic constructions, random effects, and the grand mean across them (ST).

**Figure S6.**
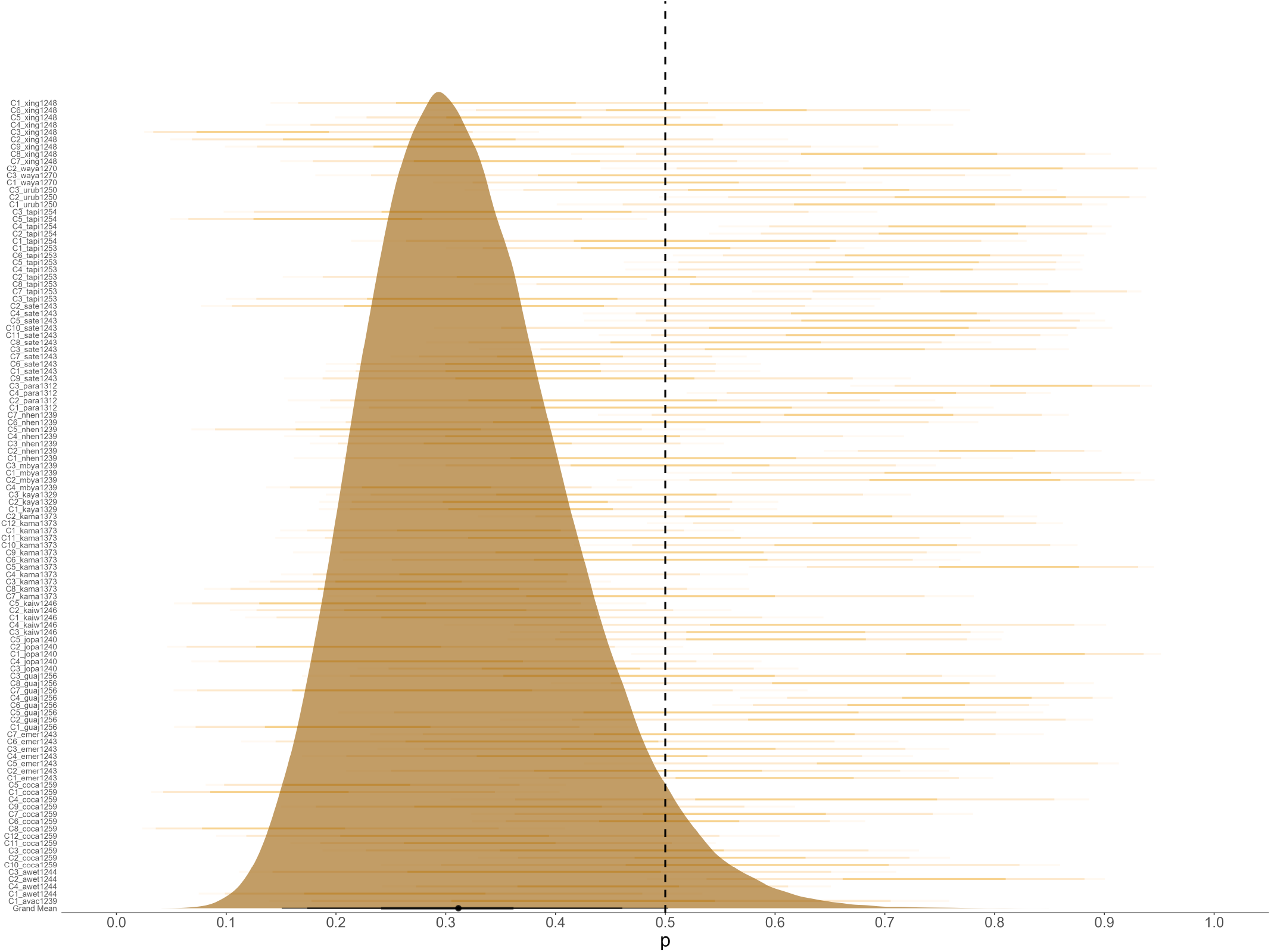
Posterior stationary probabilities (logit^™1^*θ*) for specific syntactic constructions random effects and the grand mean across them (TG).

**Table S1.** Results of Meta-Analysis I.

| Construction | Family | Median | 95% HPDI | Pr($P < .5$) |
| --- | --- | --- | --- | --- |
| ‘because’ | Indo-European | 0.18 | [0.12, 0.26] | 1.00 |
| ‘because’ | Sino-Tibetan | 0.35 | [0.25, 0.46] | 1.00 |
| ‘because’ | Tupí-Guaraní | 0.38 | [0.28, 0.49] | 0.98 |
| ‘and’ | Indo-European | 0.15 | [0.09, 0.22] | 1.00 |
| ‘and’ | Sino-Tibetan | 0.27 | [0.19, 0.38] | 1.00 |
| ‘and’ | Tupí-Guaraní | 0.31 | [0.22, 0.42] | 1.00 |
| rel. clause | Indo-European | 0.19 | [0.12, 0.28] | 1.00 |
| rel. clause | Sino-Tibetan | 0.50 | [0.39, 0.61] | 0.52 |
| rel. clause | Tupí-Guaraní | 0.46 | [0.35, 0.57] | 0.78 |

